# Learning in chunks: A model of hippocampal representations for processing temporal regularities in statistical learning

**DOI:** 10.1101/2022.04.04.487009

**Authors:** Wei Tang, Morten H. Christiansen, Zhenghan Qi

## Abstract

We investigated the neural basis of chunking during statistical learning (SL). Behavioral evidence suggests that a common mechanism in learning and memory can serve to combine smaller units into larger ones to facilitate sensory and higher-level processing. And yet, the neural underpinnings of this mechanism remain unclear. Drawing insights from previous findings of neural codes in the hippocampus, we propose a computational model to account for the temporal chunking process in SL for sequential inputs. We operationalize chunking into a hidden Markov model (HMM) that incorporates two core principles: (1) the hidden states represent serial order rather than specific visual features, and (2) the formation of temporal chunks leads to autocorrelated brain activity. We show with numeric simulations that the HMM can decode embedded triplet representations when both assumptions hold. Applying the HMM to functional neuroimaging data from subjects performing a visual SL task, we show that decoding was successful (1) for triplet sequences but not random sequences, (2) at the later stage but not earlier stage of learning, and (3) in the hippocampus but not in the early visual cortex. These results provide evidence for a hippocampal representation of generalized temporal structure emerged from sequential visual input, shedding light on the chunking mechanism for SL.

**Significance:** In statistical learning (SL), individuals develop internal representations of patterns after brief exposure to structured stimuli. People tend to recognize frequently co-occurring items as a single unit. This process, known as “chunking”, is understood to play an important role in facilitating sensory processing for learning. However, its neural underpinnings remain unclear. In this study we draw insights from hippocampal coding theories and introduce a chunking model focusing on generalized presentations for SL. With functional neuroimaging data from human subjects performing a visual learning task, the chunking model successfully decoded the temporal regularities embedded in the sequential inputs. This model and related findings provide critical evidence for a chunking process underlying SL as well as its representation in the human hippocampus.

## Introduction

In statistical learning (SL), people build internal representations of spatial and temporal regularities after brief exposure to stimuli with embedded structures. This process enables pattern segmentation from a multiplexed background or a continuous input stream, without the need for explicit instructions or feedback (Fiser & Aslin, 2001; Sherman, Graves, & Turk-Browne, 2020; Thiessen, Girard, & Erickson, 2016). For example, when exposed to streams of speech, people develop preference towards co-occurring items in their frequent order (such as English speaker’s preference for “potato” over “tapoto”). The recognition of embedded patterns becomes faster after learning, indicating that items within patterns are processed as chunks (such as the remembrance of “potato” as a whole word rather than six individual letters) (Fizer & Lengyel, 2022; Perruchet, 2006, 2019). Chunking was originally posited as a memory process which combines smaller units into larger ones to efficiently accommodate capacity limits (Halford et al., 2007; Miller, 1956; for recent considerations, see Christiansen & Chater, 2016; Thiessen 2017). In SL tasks, chunking-based models (Christiansen, 2019; Fiser & Lengyel, 2022; Orbán et al., 2008; Isbilen et al., 2022; Perruchet 2019) suggest that reappearing associative relationship among items can lead to internal representational changes that facilitate downstream cognitive processing.

Despite the behavioral models that suggest a strong link between chunking and SL, the neural basis of this link remain elusive. Theoretically, situating a chunking process in the context of SL imposes constraints on the neural representations. First, learning statistical regularities would be expected to involve a certain level of generalization, such that the internal representational changes to accommodate learned patterns should generalize (at least in part) to different input that follow the same regularity^1^. Second, chunking implies a process in which elements of a scene or input stream form an associative relationship among one another.

Previous work on generalized and associative coding during spatial navigation and memory tasks have shed important light on the potential chunking mechanisms. On one hand, neural codes in the hippocampus are found to generalize when learning to solve spatial and non-spatial tasks (Constantinescu et al., 2016; Doeller et al., 2010), leading to the theory of cognitive maps for constructing abstract knowledge (O’Keefe and Nadel, 1978; Tolman, 1948; Whittington et al., 2020). On the other hand, hippocampal neurons also code for association among objects/events that occur close in space and time (Burgess et al., 2002; Ekstrom & Ranganath, 2018; O’Reilly & Norman, 2002; Schiller et al., 2015; Schapiro et al., 2012), forming the basis of context-based learning and memory (Howard & Kahana, 2002; Sederberg, Howard, & Kahana, 2008). Recent SL studies have also found hippocampal activity sensitive to temporal associations among sequential visual stimuli (Schapiro et al., 2012, 2014; Turk-Browne, 2019), encouraging further pursuit of a chunking theory that integrates these findings.

Inspired by the previous studies, we propose a chunking model to account for the hippocampal representation in human subjects when learning embedded temporal structures from sequential inputs. We build the model on two core assumptions: *generalization* and *associative bonding*, operationalized in the form of a hidden Markov model (HMM). In the HMM, a set of hidden states define the learned internal representation, and a probability function links the hidden states to observed brain activity. To achieve *generalization*, the hidden states are set to be the serial order of items in a temporal pattern. Specifically, for embedded triplets, the hidden states are nominal variables *start*, *mid*, and *end* placed in a fixed order to represent the first, second and third positions in a triplet. This representation is invariant to the sensory features of the input items. Crucially, to relate the hidden states to brain activity, we hypothesize that (1) different hidden states correspond to distinct brain activation levels, and (2) the activity corresponding to neighboring states become similar after learning (Schapiro et al., 2012). The first assumption specifies how brain activity encodes the serial-order representation; the second assumption operationalizes *associative bonding* as a form of autocorrelated brain activity, enabling the formation of temporal chunks. We show with numerical simulations that when both assumptions hold, the HMM can decode the serial-order representation from observed data.

We tested this chunking model against hippocampal blood-oxygenation-level-dependent (BOLD) activity in subjects performing a visual SL task with an embedded triplet structure in the stimuli, while undergoing functional magnetic resonance imaging (fMRI). To test *generalization*, we used two types of stimuli (letters & pictures) presented in intermittent blocks. In each learning session, a triplet structure was embedded in one type of the stimuli while the other type was presented in a random order. We used this setup to test whether the hippocampal BOLD activity was representative of serial order regardless of stimulus type. To test *associative bonding*, we compared the decoding performance of two HMMs, one taking into account the autocorrelation of BOLD activity between neighboring stimuli while the other one did not. The decoding efficacy of the two models would reveal the validity of the underlying model assumptions. We were aware that multiple sources could contribute to the autocorrelation of the BOLD signal, including noncognitive, metabolic processes unrelated to the hypothesized neural representation. Accordingly, we used the simulation to understand how chunking-related and unrelated autocorrelation would affect the decoding performance, which informed our interpretation of the empirical BOLD decoding results. Our findings provide further insight into the chunking process for SL and its representation in the hippocampus.

## Results

### Overview

We used a classical triplet SL paradigm to test the chunking model. The task stimuli contained series of images presented one at a time, half of which were embedded with a triplet structure and the other half randomly ordered (Fig. 1A). We tested how the human brain represented the embedded triplets with a temporal chunking model. The model took the form of an HMM, which used a probabilistic function (“emission probability”) to link the hypothesized internal representation and the observed brain activity. The rationale was that if the model assumptions were biologically plausible, then the HMM would be able to decode the triplet representation from the observed brain activity when the triplet series were presented. Because our hypotheses were drawn from findings of the hippocampus, we particularly tested the decoding accuracy with hippocampal activity. We note that this chunking model focused on the neural representation rather than a specific learning algorithm that led to the representation. Nonetheless, the HMM approach allowed us to identify whether and when the hypothesized representation emerged as an outcome of SL.

**Figure 1.**
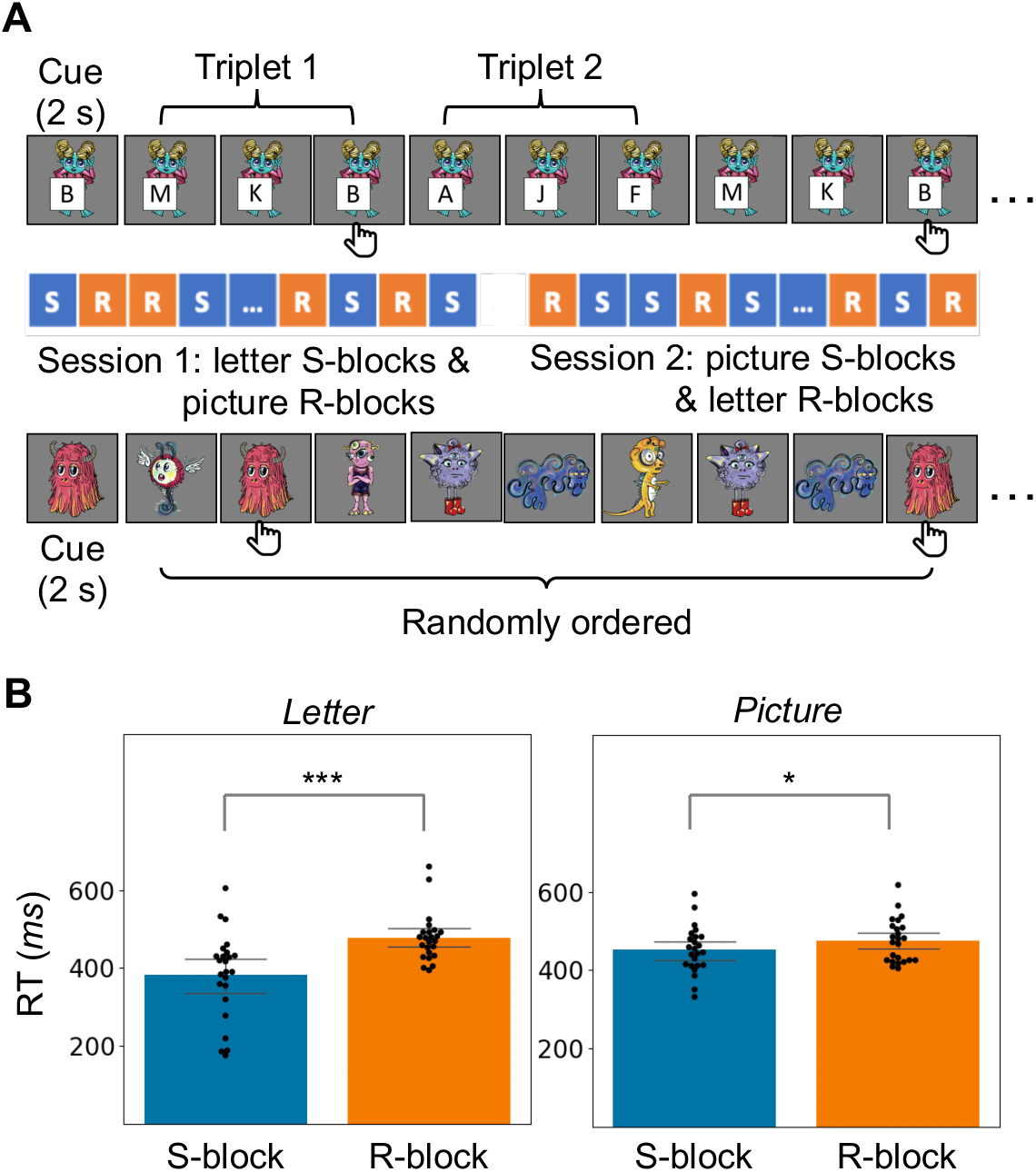
Behavioral sensitivity to temporal structures in sequentially presented visual stimuli. (A) Schematic of task design. Dashed vertical line separates two learning sessions, each composed of 6 S-blocks (blue) intermixed randomly with 6 R-blocks (orange). Each mini block started with a cue for the target, followed by 48 stimuli presented sequentially. Subjects responded to the target with a button press. (B) RT of the structured and random sequences for each type of stimuli, shown as median ± s.e. across subjects. Black dots on the swarm plot represent the cross-trial median of individual subjects. Asterisks mark the significance of the Wilcoxon test, ***: *p* < 0.001, *: *p* < 0.05.

### Behavioral learning effect in the visual SL task

Human adult participants viewed sequentially presented images while responding to target images embedded in each sequence (Fig. 1). The sequence blocks varied along two dimensions: the stimuli in each block were (1) *letters* or *pictures* and (2) temporally arranged into triplets (S-block) or presented in random order (R-block). The temporal structure was set to systematically differ between *letter* and *picture* stimuli across two learning sessions (Fig. 1A). In Session 1, *letter* S-blocks were pseudo-randomly intermixed with *picture* R-blocks. In Session 2, the structure/randomness was switched, such that *letter* R-blocks were pseudo-randomly intermixed with *picture* S-blocks. The target was set to be the third element of a randomly selected triplet from either the *letter* or *picture* sequences, and its location followed no systematic pattern in each block either within– or across-participants. Subjects were not informed of the temporal structure and completed the target-detection task with an average hit rate of 97.4 ± 4.7% (cross-subject median ± s.d.), false-alarm rate of 0.2 ± 0.2%, and reaction time (RT) of 446 ± 52 ms. Comparisons of RT between S– and R-blocks showed a facilitation effect on target detection. For both types of stimuli, across subjects, the trial-average RT was significantly lower in S-blocks than in R-blocks (Fig. 1B; *letter* stimuli: Wilcoxon signed-rank *W* = 23.0, *p* < 0.00028; *picture* stimuli: *W* =71.0, *p* < 0.024).

### Theoretical construct of the chunking model

To model the brain representation of the triplet structure embedded in the stimuli, we made two core assumptions in building the HMM: *generalization* and *associative bonding*. The HMM contained three nominal hidden states: *start*, *mid* and *end*, following the rationale that all triplets in the input contained a start, a middle and an end position regardless of the sensory content at each position. By construction, the transition among hidden states in a triplet sequence was only possible from *start* to *mid*, *mid* to *end*, and *end* to *start* (Fig. 2A). Together, the hidden states and their transition pattern formed a generalized, serial-order representation of embedded triplets. The hidden states were linked to brain activity via an emission probability function (*b*(*o_t_*) in Fig. 2A), which specified how the three states drove the BOLD activation into three distinct levels. The separation between levels was controlled by a parameter *d’* in the emission probability. To operationalize *associative bonding*, we let the BOLD activation for a hidden state be informative of the BOLD activation for its neighboring states in a sequence. This associative relationship manifested as an autocorrelation parameter *ρ* in the emission probability. With *d’* and *ρ*, the model established a probabilistic relationship between the time-to-time BOLD fluctuation and hidden state sequences (Fig. 2B). For brevity, we refer to this model as the cHMM (c for “chunking”).

**Figure 2.**
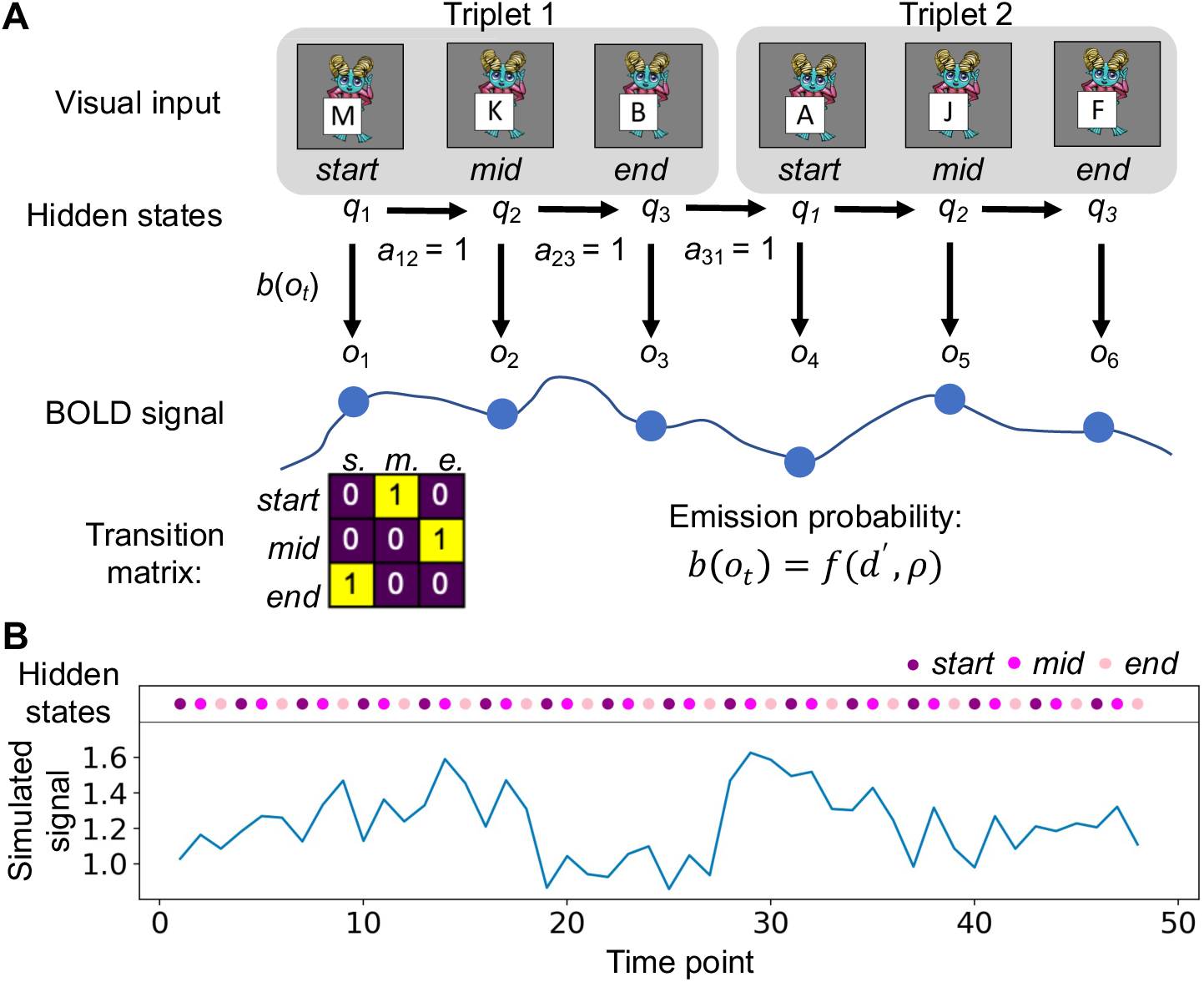
The HMM design. (A) Schematic of the HMM for testing the chunking hypothesis. Three hidden states *start*, *mid*, and *end* were defined according to the position of each stimulus in a triplet, regardless of their specific visual content. The triplet structure was defined by the transition matrix (abbreviations: s. for *start*, m. for *mid,* e. for *end*), in which the transitions from *start* to *mid*, *mid* to *end*, and *end* to *start* had probabilities of 1. Each hidden state was linked to the observed BOLD signal by the emission probability, which followed a conditional-Gaussian distribution. (B) An example of simulated time series from the cHMM, with parameters *d’* = 0.9, ρ = 0.4.

To test model validity, we devised a decoding analysis which attempted to estimate the hidden-state transition matrix from observed brain activity. The rationale was that the cHMM made specific assumptions of how the observed data and hidden states should relate, and accordingly, violation to these assumptions would corrupt decoding accuracy. We took note of a potential confounding factor in this approach: the autocorrelation of the BOLD signal that the cHMM used to estimate *ρ* might arise from noncognitive, metabolic processes unrelated to the neural representation being studied. To tease apart chunking-induced and other effects in the autocorrelation, we built a separate model referred to as the ncHMM (nc for “no chunking”), whose emission probability depended only on *d’*. Conceptually, if the brain activity indeed encoded serial order but not the associative relationship among states, then the information from *d’* alone would be sufficient for both the ncHMM and the cHMM to decode the hidden-state transitions. If the brain encoded both the serial order and the state association, then the cHMM would be more powerful than the ncHMM in decoding, given the cHMM’s ability to utilize information from both *d’* and *ρ*. If the assumption on serial-order representation was violated, or if the confounding effects in the autocorrelation were overwhelming, both models would fail to decode. Thus, by comparing model performance, we would gain insights into the mechanisms that generated the data. Before applying this approach to the experimental fMRI data, we tested its validity using simulated time series from different emission probability functions.

### Simulation results

We generated time series from a 48-state sequence containing 16 triplets of serially ordered *start*, *mid* and *end* states (this setup was to match the behavioral task structure; see Methods). The same sequence generated five different datasets following five emission probability functions. The function for dataset 1 had high *d’* and *ρ* values, referred to as a “chunking code”; the function for dataset 2 did not code for association between items (i.e., *ρ* = 0), and thus referred to as a “position-only code”. Both datasets contained unrealistically low noise (i.e., high *d’*), and accordingly, a strong serial-order representation that was unlikely to find with empirical BOLD data. Indeed, for the two examples shown in Fig. 3A & 3B, *d’* alone provided sufficient information for decoding, and both cHMM and ncHMM recovered the transition matrix accurately. In comparison, dataset 3 was generated with low *d’* and *ρ* values in the emission probability function, referred to as a “noisy chunking code”; dataset 4 was generated with low *d’* and *ρ* = 0, a “noisy position-only code”. The ncHMM failed to decode both examples in Fig. 3C & 3D, suggesting that information from *d’* alone was insufficient for decoding either dataset. By contrast, the cHMM decoded the example from dataset 3 accurately (Fig. 3C), demonstrating the power of utilizing both *d’* and *π*. With *π* unavailable in dataset 4, the cHMM also failed to decode (Fig. 3D). Finally, when there was no serial-order representation in dataset 5 (i.e., *d’* = 0 and *ρ ≠* 0, mimicking activity with chunking-unrelated autocorrelation), both models failed to decode (Fig. 3E). Indeed, when we disrupted serial-order information by temporally permuting each dataset, both cHMM and ncHMM failed to decode likewise (Fig. 3A-E, lower panels). We note that the stereotypical pattern of the negative cHMM outcome was an inevitable result from our decoding technique. See Methods for details.

**Figure 3.**
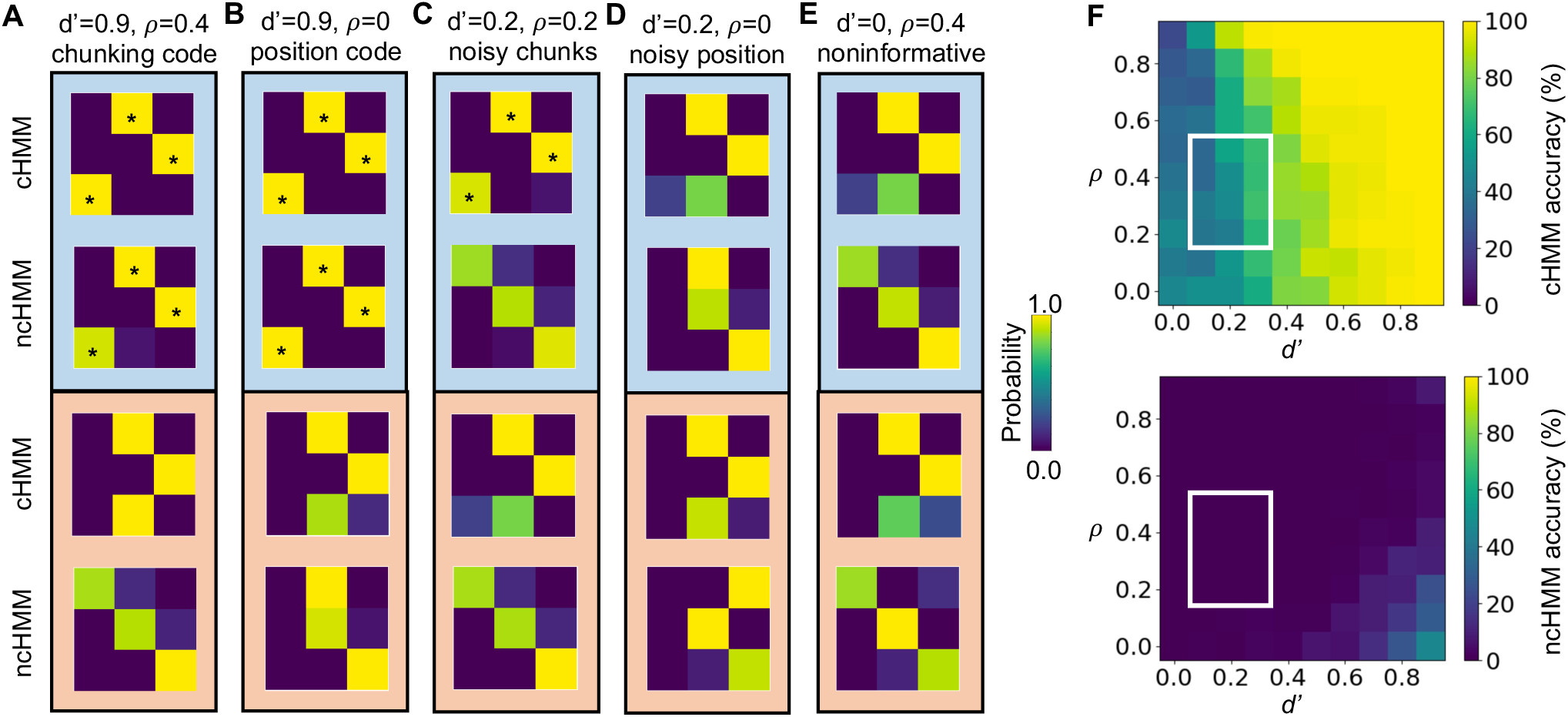
Decoding triplet representation from the simulated data. The chunking (cHMM) and no-chunking (ncHMM) were applied to 5 datasets generated with (A) a chunking code, (B) a position-only code, (C) a noisy chunking code, (D) a noisy position-only code, and (E) a null code without triplet representation. The decoded transition matrix is shown for each original dataset (blue background) and the corresponding time-permuted datasets (orange background). To facilitate comparison with the ground truth, asterisks were added to matrices that followed the cardinal pattern in Fig. 2A (i.e. the highest probabilities were found for the *start-mid*, *mid-end* and *end-start* pairs). (F) Decoding accuracy across parameter ranges. White contour marks the physiologically realistic ranges of *d’* and *ρ* calculated from the current experimental BOLD data.

The above examples illustrated the model performance on data generated from particular *d’* and *ρ* values. To gain a more comprehensive view, we varied *d’* and *ρ* systematically and examined the decoding accuracy as a function of the two variables (Fig. 3F). The accuracy was measured by the percentage of samples (100 for each parameter set) whose decoded matrices showed the same cardinal pattern as that of the ground-truth matrix (Fig. 2A). Across all the tested parameter sets, the cHMM showed overall above-chance level accuracy (chance level=1.19%, see Methods), while the ncHMM surpassed the chance level only when *d’* was above 0.5 and *ρ* was below 0.5. Within the parameter regime of the veridical BOLD data (median ± inter-quartile interval across subjects: *d’* = 0.21 ± 0.1, *ρ* = 0.39 ± 0.15, marked as white contour in Fig. 3F), the cHMM accuracy was well above-chance (within-contour median ± inter-quartile interval across parameter sets: 0.62 ± 0.14) while the ncHMM accuracy stayed at 0 (0.00 ± 0.00).

### Learning effect in the autocorrelation of the BOLD activity

A core assumption of the chunking model was the *associative bonding* between items within the triplet. This assumption predicted that brain activity would become more autocorrelated as a result of triplet representation. Indeed, for both types of stimuli, we observed significantly greater BOLD lag-1 autocorrelation across hippocampal voxels in S-than in R-blocks (Fig. 4A; *letter*: paired-sample *t*_99_ *=* 83.58, *p* < 1× 10^-6^; *picture*: paired-sample *t*_99_ *=* 77.12, *p* < 1× 10^-6^). To further test the learning effect in the BOLD autocorrelation, we examined the S-R difference of BOLD autocorrelation across mini blocks and compared its pattern with the pattern of RT. By design, the same type of stimuli switched from structured to random or vice versa between the two learning sessions. Accordingly, we expected the BOLD autocorrelation and RT to follow this change. For both metrics, we subtracted R-blocks from S-blocks for Session 1 and S-blocks from R-blocks for Session 2. This subtraction order was to facilitate the detection of a sign-flip: Under the generalized serial order assumption, the detailed visual features would not affect how the triplet was represented. Thus, the difference scores should be driven by the S/R contrast, which flipped the sign between sessions, rather than by the stimulus-type contrast, which maintained the sign (*letter – picture* for both sessions). Indeed, the sign-flip was evident in both RT and the autocorrelation of voxel-averaged hippocampal BOLD signal (Fig. 4B, within-subject main effect of session, RT: *F*_1, 19_ = 21.44, *p* < 0.00018; autocorrelation: *F*_1, 82.30_ = 9.00, *p* < 0.0035). Moreover, the RT difference and the autocorrelation difference, averaged across participants, were correlated across mini blocks (Fig. 4C, Spearman’s *r=* –0.62, *p <* 0.035), such that a greater reduction in RT corresponded to a greater increase in the BOLD autocorrelation.

**Figure 4.**
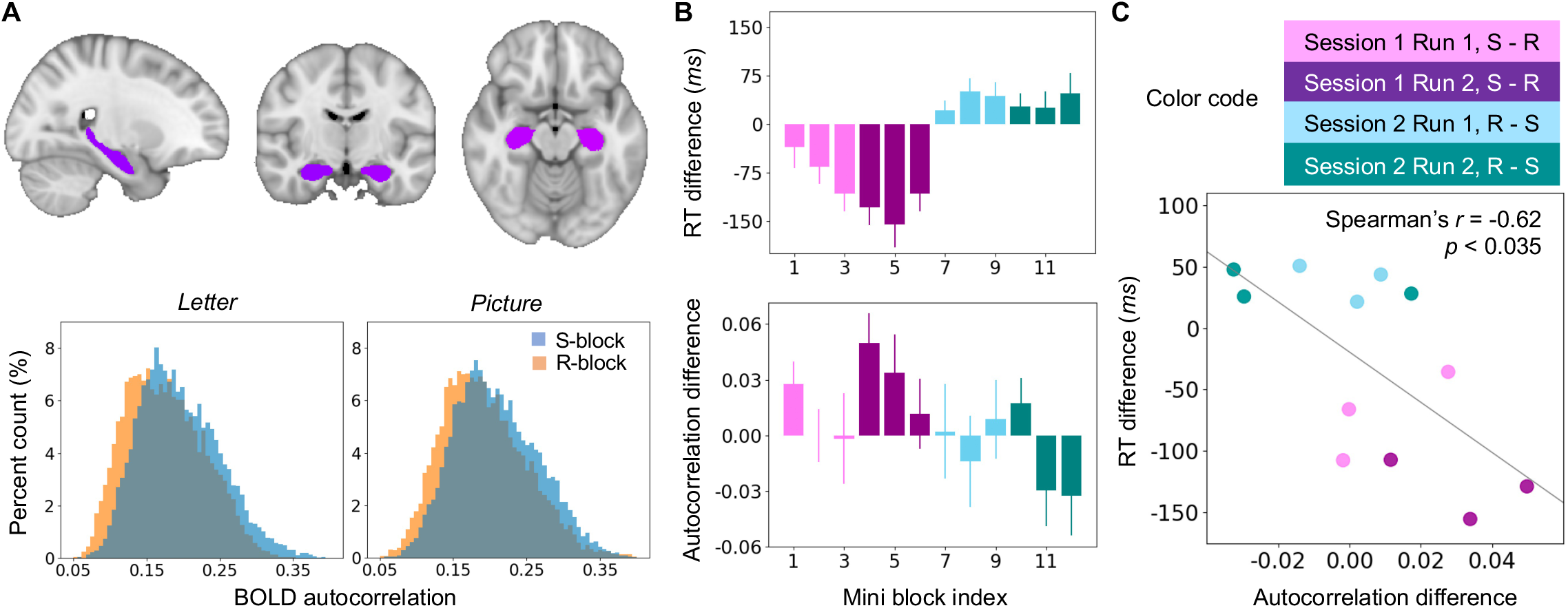
BOLD autocorrelation sensitivity to the structured stimuli. (A) Voxel histograms (bottom) of subject-averaged BOLD autocorrelation in the hippocampus extracted from the MNI152 template (top, purple color marking the hippocampus). The BOLD autocorrelation from each subject’s native MRI space was first registered to the template space and then averaged within-voxel. (B) The average S-R difference of RT and BOLD autocorrelation across mini blocks, shown as median ± s.e. across subjects. (C) The correlation across mini blocks between the S-R difference of RT and the S-R difference of autocorrelation. Each dot shows an averaged value across subjects.

### HMM decoding with hippocampal BOLD data

We applied the cHMM and ncHMM to decode triplet representations from the hippocampal activity (Fig. 5). Both analyses were performed on the voxel-averaged BOLD time series. We examined the decoding accuracy at the earlier and later stage of learning by analyzing the two runs of each session separately. The cHMM was able to decode the triplet representation in the second but not the first run of each session, from the S-blocks but not the R-blocks. This decoding pattern was consistent for *letter* and *picture* stimuli (Fig. 5A). In contrast, the ncHMM failed to decode the triplet pattern in either session, from either type of block, or for either type of stimuli (Fig. 5B). As a control test for potential false-positive decoding caused by the BOLD autocorrelation unrelated with chunking, we also applied the cHMM to the voxel-averaged BOLD time series from the primary visual cortex (Fig. 5C). The simulation suggested that signals generated from high *ρ* but no serial-order representation (*d’* = 0) would fail the decoding. We estimated *d’* and *ρ* using the mean and autocorrelation values of BOLD activity in the hippocampus and V1. The V1 activity showed much lower *d’* (*t* = 11.09, *p* < 9.58 × 10^-10^) and higher *ρ* (*t* = –11.10, *p* < 9.49 × 10^-10^) than the hippocampal activity (Fig. 5D). No triplet representation was decoded from V1 using either session, either type of block, or either type of stimuli (Fig. 5E).

**Figure 5.**
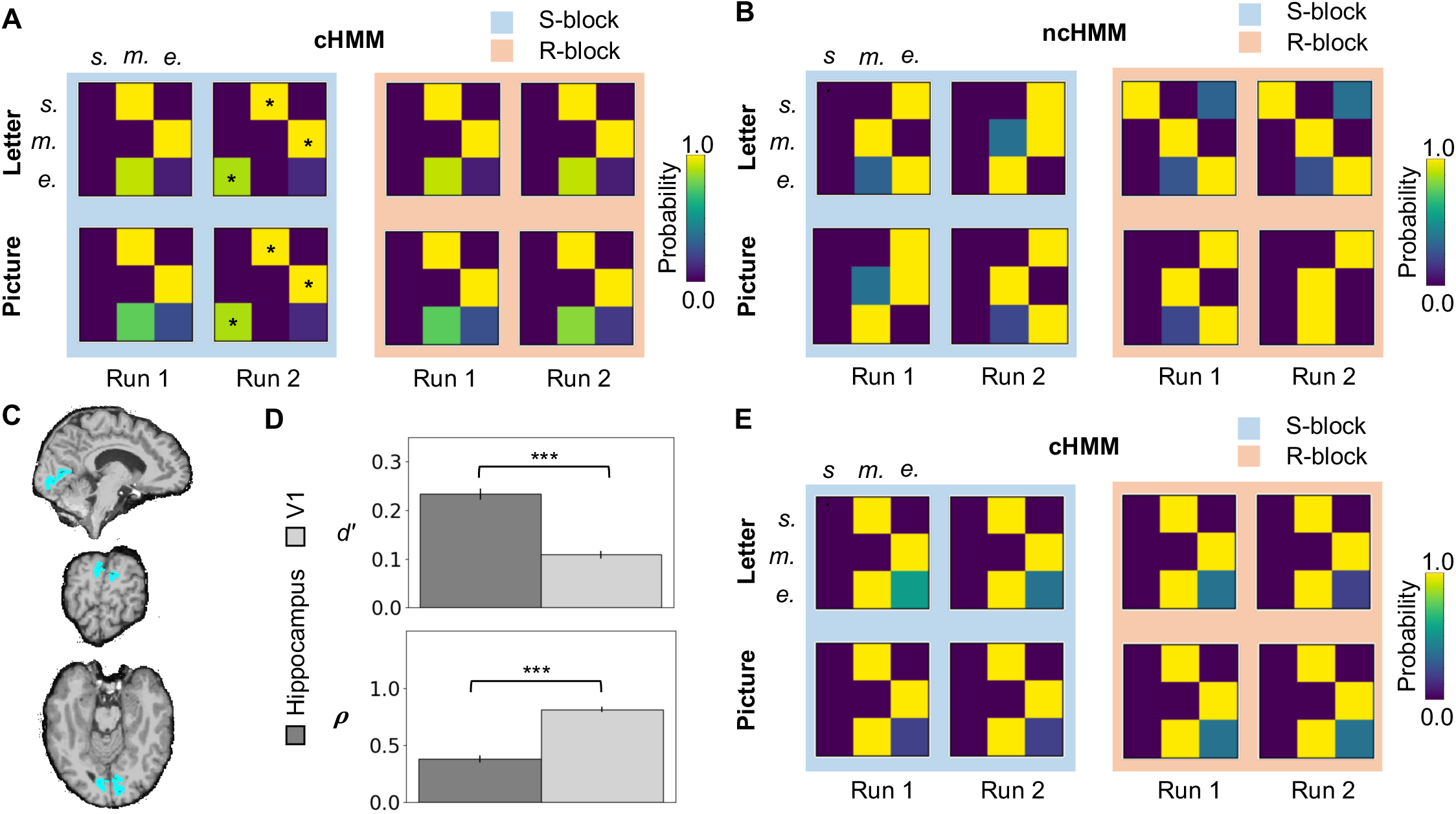
Decoding triplet representation with the veridical BOLD activity. (A) The cHMM and (B) the ncHMM decoded transition matrices from the hippocampal activity. Asterisks were added to matrices that resembled the ground-truth transition matrix. (C) The mask of V1 region shown in an example subject’s brain. (D) Estimated *d’* and *ρ* of the hippocampal and V1 activity, shown as the mean ± s.e. across subjects. Asterisks mark the significance of the paired-sample *t* test across subjects, ***: *p* < 0.001. (E) The cHMM-decoded transition matrices from the V1 activity.

## Discussion

We propose a chunking model for SL based on generalized and associative representations of sequential inputs. This chunking model makes two core assumptions: (1) the generalized representation of serial order leads to distinct brain activation levels (measured by *d’*), and (2) an *associative bonding* mechanism leads to structured autocorrelation in the brain activity (measured by *ρ*). We operationalized this theory into a chunking HMM and tested its capability in decoding triplet structures from the hippocampal activity, using fMRI data of subjects performing a visual SL task.

The HMM approach provides a principled framework for testing hypothesized internal representations in the observed brain activity. Generally, HMMs deploy probability theory to infer hidden states based on trial-to-trial variation in the observed data. In the current study, the chunking hypothesis is built into the emission probability function linking the hidden states to *d’* and *ρ* that characterize the BOLD measurements. The particular form of this function determines how a triplet representation drives the BOLD data. Crucially, when the emission probability depends on both *d’* and *ρ*, it assumes not only that the serial order drives the mean BOLD fluctuation, but also that the associative relationship between neighboring states leads to BOLD autocorrelation. This emission probability provides an operational definition of the chunking process. The numerical simulations showed that if both assumptions hold, the cHMM can decode the hidden states from the observed data. In the parameter regime that simulated the veridical BOLD characteristics, i.e. low *d’* and moderate *ρ* (Fig. 3F), the cHMM decoded well above chance level by taking into account both *d’* and *ρ* information, while the ncHMM failed to decode by relying only on *d’*.

These principles guided the interpretation of the hippocampal BOLD decoding results. We found that (1) the cHMM decoded the triplet representation from the hippocampal activity in the S-blocks but not the R-blocks, for both types of stimuli, and (2) the ncHMM failed to decode any triplet representation from either type of stimuli. The insensitivity to stimulus type was evidence of generalization, and the cHMM success versus ncHMM failure validated the role of BOLD autocorrelation in triplet representation. Moreover, the cHMM decoding was successful only in the later stage (second run) of each session, indicating that the triplet representation was gradually formed upon repeated exposure. The independent analysis of BOLD autocorrelation supported these insights. First, the hippocampal BOLD autocorrelation was greater in S-blocks than in R-blocks regardless of stimulus type. Second, the increase was proportional to the reduction of RT, establishing a link between hippocampal activity and the behavioral effect of facilitated target detection. Taken together, the HMM and autocorrelation analyses cross-validate each other in supporting the two assumptions of the chunking model. The hippocampal representation is in a generalized form sensitive to serial order rather than stimulus details, and it involves a binding mechanism manifested as increased BOLD autocorrelation upon repeated exposure to structured input.

Importantly, the decoding result suggested against the scenario that the hippocampus represented item association alone. One could argue that chunking can be achieved by associating items together without a generalized representation of serial order. However, the numerical simulation suggested against this scenario. The cHMM performed poorly when *d’* was 0 even if *ρ* was high (dataset 5). The decoding result for V1 further verified this point. The BOLD signal of V1 showed significantly greater *ρ* and lower *d’* than that of the hippocampus. Nonetheless, this high *ρ* value was insufficient to inform the triplet representation, and the cHMM failed to decode in either session with either type of stimuli. In fact, V1 neurons are known to process lower-level visual features rather than forming generalized representations. This result also helped ruling out the possibility that the hippocampal decoding was an artifact driven by metabolically generated autocorrelation rather than task-related autocorrelation.

This chunking model draws inspiration from long-standing hippocampal coding theories. Studies of spatial navigation and long-term memory have observed different forms of generalized representations that lead to the theory of cognitive maps (Tolman, 1948; O’Keefe et al., 1978). In a separate endeavor, studies of the hippocampal responses to objects/events occurring close in time or space have suggested associative coding (Burgess et al., 2002; Cohen & Eichenbaum, 1995; Ekstrom & Ranganath, 20), which is at the core of context-based memory (Howard & Kahana, 2002; Sederberg, Howard, & Kahana, 2008). Although the two lines of research focused on different aspects of hippocampal activity, with distinct viewpoints yet to be reconciled (Lisman et al., 2017), recent models have suggested that generalized and associative coding can be complementary in assisting the process of learning (Flesch et al., 2023; Whittington et al., 2020). Experimentally, a recent SL study with human intracranial recording has found serial order representation among the hippocampal neurons (Henin et al., 2021). Such theoretical and empirical evidence encouraged us to explore *generalization* and *associative bonding* as the basis of chunking. Although the current model does not specifically address how the chunks are learned, it nonetheless allows us to examine when such representations arise in the brain at different stages of SL. To further develop a learning mechanism, future work is needed to determine how more general constraints from stimulus– and modality specificity can be incorporated and explored empirically (see e.g., Frost et al., 2015, 2019, for discussion). The current model also has its potential to connect with context-based models for memory retrieval and future prediction (e.g., inference of previous/future hidden states based on the learned associative relationship in the emission probability function). Further development in this direction will help elucidate the shared mechanisms underlying implicit learning, memory and prediction (Reber, 2013; Sherman and Turk-Browne, 2020), while allowing empirical tests with human *in vivo* data.

Finally, we note that the HMM framework is versatile for testing different learning models on a variety of data. The utility of HMM is two-fold: with a fixed form of emission probability, one can compare different theories on the internal representation by manipulating the hidden states; alternatively, with a fixed form of internal representation, one can explore different brain mechanisms that generate the observed activity by manipulating the emission probability. The current work takes the second path with one specific chunking hypothesis, while ample options are available for future exploration. For example, one can employ more sophisticated models to set up the hidden states, e.g., those following the hierarchical Bayesian theory (Orbán et al., 2008). The emission probability can be extended to a multivariate form to account for multi-voxel BOLD activity. The emission probability can also be fit to different types of data including invasive neural recordings.

## Methods

### Participants

Twenty-eight adults (mean age = 20.79 years, SD = 2.89 years, 7 males) participated in this study. All participants were right-handed, native English speakers with no history of neurological or psychiatric disorders. All participants gave written consent to perform in accordance with the Institutional Review Board at the University of Delaware. All participants were compensated for their participation. Due to incomplete MRI data resulting from acquisition issues (N = 2), early termination of the study by the participants (N = 2), poor performance with < 20% hit rate in at least one scanning session (N = 4), twenty participants in total (mean age = 20.16 years, SD = 0.81 years, 7 males) were included in the data analyses.

### Stimuli

The stimuli contained two types of images: *Letters* (linguistic stimuli) were 12 images each containing an alien character holding up a sign with a capital letter (see Supplemental Figure for all the stimuli). The alien character was held the same while the letter differed among the 12 images. *Pictures* (nonlinguistic stimuli) were 12 different standalone alien cartoons (Supplemental Figure). Both Letters and Pictures were presented in two types of mini blocks. An *S-block* was composed of 48 stimuli, organized into 4 unique triplets each repeated 4 times. The presentation order was pseudo-randomized among triplets, so that the TP between stimuli was 1 within-triplet and 1/4 between-triplet. No *letter* triplet contained any words, common acronyms, or initialisms. An R-block contained 12 unique stimuli each repeated 4 times. In other words, an R-block was the randomization of all stimuli in an S-block of the same stimulus type. The presentation order of individual stimuli was pseudo-randomized, and no combination of three stimuli was presented more than once. Thus, the TP among stimuli was uniformly 1/12. For both S– and R-blocks, each stimulus was presented at the center of the screen for 0.8 s with an 0.2 s interval between stimuli. Each mini block lasted 48 s.

### Procedure

Participants watched two sessions of stimuli presentation, organized into mixed S– and R-blocks described above, while performing a target detection task. Session 1 contained 6 linguistic S-blocks, 6 nonlinguistic R-blocks and 6 12-second resting periods; Session 2 contained 6 nonlinguistic S-blocks, 6 linguistic R-blocks and 6 12-second resting periods. The order of S-blocks, R-blocks and resting periods was pseudo-randomized, and the randomization was independent for each session and each participant. Adjustment was made to ensure that no more than two blocks of the same type were adjacent to each other. Before each block, a cue for the target was presented at the center of the screen for 2 s. For both linguistic and nonlinguistic stimuli, the cue was a stimulus chosen from the last position of a triplet in an S-block. The cue was randomly assigned from the four triplet options for each participant. Throughout the task, in each participant and for each type of stimuli, the same cue was used for all S– and R-blocks. Because the order of stimuli and that of blocks were randomized independently for each participant, the target location among stimuli followed no systematic pattern either within– or cross-participants.

### The HMMs

The HMM modeling followed the rational that all the triplets contained a start, a middle and an end position regardless of the sensory content at each position. Accordingly, the HMMs contained 3 hidden states:

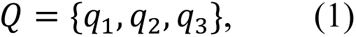

where *q*_1_, *q*_2_, *q*_3_ denote the first (*start*), second (*mid*) and third (*end*) positions in a triplet. The triplet representation would be the three states placed in a serial order, *start-mid-end*. The transition probability among hidden states was defined by the matrix

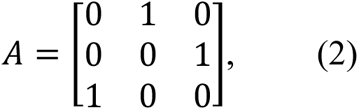

Each element on the *i-*th row and *j*-th column represented the probability of transition from *q_i_* to *q_j_*. By construction, the transition for a triplet sequence was only possible from *start* to *mid*, *mid* to *end*, and *end* to *start*. Thus, *A* had a probability of 1 in the corresponding cells and 0 elsewhere. The initial probability of *Q* was set to *π* = [1, 0, 0], because the triplet sequences in this study all began with *start*.

The hidden states were linked to brain activity via an emission probability function. Denote the simulated or experimental BOLD time series as *O*(*t*). The relationship between the hidden states and *O*(*t*) was determined by *b*(*o_t_*|*q_j_*), the probability of observing activity *o* at time *t* given state *j*. Critically, *b*(*o_t_*|*q_j_*) was hypothesized to follow a conditional Gaussian distribution:

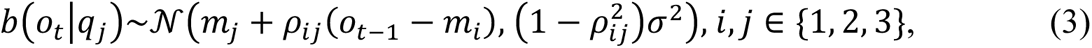

where *m_i_* and *m_j_* denoted the mean BOLD activity corresponding to hidden states *i* and *j*, respectively, *ρ_ij_* the correlation of the BOLD activity corresponding to hidden states *i* and *j*, and *σ* the standard deviation that was assumed to be constant in the BOLD time series. The right-hand side of Eq. 3 was the conditional distribution of *o_t_* (corresponding to state *j*) given *o_t_*_-1_ (corresponding to state *i*), assuming a bivariate Gaussian joint distribution. Note that *o_t_* and *o_t_*_-1_ denoted measurements at adjacent time points, and therefore their correlation *ρ_ij_* was the lag-1 autocorrelation of the BOLD activity. Eq. 3 stated that given hidden state *j*, the probability of observing *o_t_* was a function of state-dependent variables *m_i_*, *m_j_* and *ρ_ij_*. For triplet sequences, because the transitions among states were deterministic, the *i-j* pairs were unique, i.e., there was a unique *m_i_* for each *m_j_*. Thus, *b*(*o_t_*|*q_j_*) was unique for each state *j* at time *t*. For generating simulated data (see *Simulation*), *ρ_ij_* was set equal for all *i-j* pairs; for the decoding analysis (see *Decoding triplet representation*), *ρ_ij_* was estimated separately for each *i-j* pair from observed data. In describing the simulation and empirical results, we reported the average *ρ_ij_* across state pairs as a single-valued *ρ* for conformity.

### Triplet representation with and without the chunking code

The presence of autocorrelation *ρ_ij_* in *b*(*o_t_*|*q_j_*) set apart our model from previous HMMs that used a univariate Gaussian emission probability to detect task-related representation patterns in fMRI data (Baldassano et al., 2017; Duan & Man, 2012). In a univariate-Gaussian framework, the setup for hidden states *Q*, transition matrix *A* and initial states *π* would be the same as above, but the emission probability would depend only on the mean and variance of the BOLD signal at time *t*:

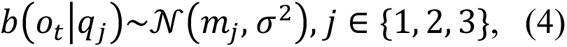

In Eq. 4, the only state-dependent parameter was the mean BOLD activity *m_j_*. This emission probability assumed that different hidden states result in different mean levels of BOLD activity, controlled by the sensitivity index *d’*:

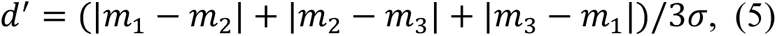

The value of *d’* would determine how well the observations corresponding to the three states can be separated by their mean values. A high value of *d’* would correspond to a clear mean separation and hence a strong representation of serial order. This univariate-Gaussian HMM could achieve a generalized representation of triplets with the serial order assumption, but there would be no representation for the associative relationship among states.

In comparison, the conditional-Gaussian HMM could achieve *associative bonding* with the autocorrelation parameter *ρ_ij_* in Eq. 3. A nonzero *ρ_ij_* would allow the activity evoked by the previous state to influence the activity of the current state. A high value of *ρ_ij_* would correspond to high predictability from state to state. Notably, when *ρ_ij_* would become zero for all state pairs, Eq. 3 would become Eq. 4 and the model would degenerate to a univariate-Gaussian HMM. With both *d’* and *ρ_ij_* being nonzero, the conditional-Gaussian HMM would represent triplets with both the generalized serial order and the state bonding, meeting our two assumptions for the chunking mechanism in SL. We referred to the conditional-Gaussian model as cHMM (c for “chunking”) and the univariate-Gaussian model as ncHMM (nc for “no chunking”).

### Decoding triplet representation

Model validity was tested via a decoding procedure using the Baum-Welch algorithm (Jurafsky & Martin, 2009), in which the hidden-state transition matrix *A* was treated as an unknown parameter to be estimated, or decoded, from the observed data. To initiate the Baum-Welch algorithm, an uninformative prior of *A* was set up such that each probability in the matrix was 1/3. The priors of parameters in *b*(·) were estimated from the time series to be decoded. Each simulated or empirical BOLD time series *O*(*t*) contained 48 data points corresponding to the 48 stimuli (see *Stimuli* section below):

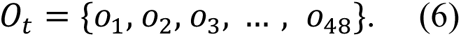

The initial values of parameters in Eqs. 3 & 4 were calculated from *O*(*t*):

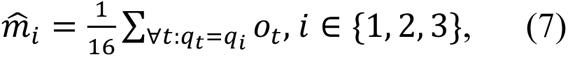

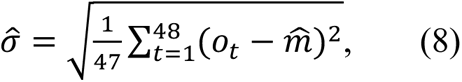

where 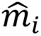 denoted the mean of *o_t_* across time points corresponding to state *i*, and 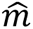 the mean across all time points. The initial value of *ρ_ij_* was calculated separately for each *i-j* pair:

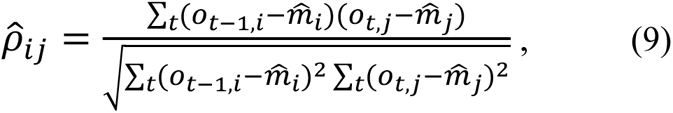

in which the summation was over all time points associated with state pair *i-j.* With the initial estimate of *A* and *b*(·) and fixing *π* to [1, 0, 0], the Baum-Welch algorithm would re-estimate *A* and *b*(·) recursively through an expectation-maximization procedure until the results converged. The decision to force *π* = [1, 0, 0] rather than estimating it from the data was due to the limited sample size for estimation. The full sequence of both the simulated and experimental data contained only 48 data points, a moderate size for estimating transition patterns among three states. To reduce the number of unknown parameters, we considered *π* = [1, 0, 0] a reasonable choice because all the tested triplet sequences began with *start*. For the decoded matrix to reveal a triplet transition pattern, the highest probability values would need to be attributed to the *start-to-mid*, *mid-to-end* and *end-to-start* transitions. In other words, the decoded matrix had to follow Eq. 2 in a cardinal sense if the highest and lowest values were not ones and zeros.

### Decoding outcome for randomly ordered sequences

As a technical note, the cHMM decoding output for random sequences would have a stereotypical pattern, such that the highest transition probabilities fell on *start-to-mid*, *mid-to-end* and *end-to-mid* if the data contained no triplet representation. This was a theoretical result due to the way priors were set up. Essentially, in the first few rounds of the Baum-Welch algorithm, because the initial probability was fixed to *π* = [1, 0, 0], and because *b*(·) was initialized from data mean and autocorrelation, the initial maximum-likelihood estimate of the state sequence would almost^2^ always begin with the three states *start*, *mid* and *end* in such order. The initial noninformative transition matrix *A* would predict the upcoming states to be uniformly random among *start*, *mid* and *end*. Based on the predicted sequence, the iterative algorithm would then re-estimate *A* for the next round of prediction. Because the predicted sequence was largely random, the first three states would nudge the re-estimated *A* to have slightly higher probabilities for *start-to-mid* and *mid-to-end* transitions. This bias would affect state prediction in the second round: Following *end*, if the next state was *start*, the bias in *A* would make *mid*-*end* the most probable upcoming states; if the next state was *mid*, then the most probable subsequent state would be *end*; if the next state was *end*, then the subsequent state would be *start/mid/end* with equal probability. The periodicity of these patterns suggested that state pairs *mid*-*end* and *end-mid* would appear the most frequently in the predicted sequence, resulting in higher re-estimated probabilities of *mid-to-end* and *end-to-mid* in *A* for the next round. Through iteration, the favored transitions would reinforce one another, resulting in the stereotypical pattern in the final output. Compared to the cHMM, the ncHMM had more variability in initializing the first three states, because it required only the mean values to initialize *b*(·). Thus, the ncHMM would allow more flexible transition patterns in *A*.

### Simulation

Stochastic processes were generated from the HMMs for testing decoding efficacy. HMMs are generative models, i.e. given *π*, *A*, and *b*(·), samples of O(*t*) can be drawn from the emission probability distribution up to an arbitrary time point *T.* To match the task setup, *T* was set to 48 for all simulated time series. Five sets of simulated data were generated with (1) *d’* = 0.9 and *ρ* = 0.4, corresponding to the strong encoding of serial order and *associative bonding* (i.e., a “chunking code”), (2) *d’* = 0.9 and *ρ* = 0, corresponding to the encoding of serial order without *associative bonding* (“position-only code”), (3) *d’* = 0.2 and *ρ* = 0.2, corresponding to the physiologically realistic weak encoding of serial order and *associative bonding* (“noisy chunking code”), (4) *d’* = 0.2 and *ρ* = 0, corresponding to the physiologically realistic weak encoding of serial order without *associative bonding* (“noisy position-only code”), and (5) *d’* = 0 and *ρ* = 0.4, corresponding to an autocorrelated signal with no triplet representation (“null code”). Dataset 5 was to simulate the ubiquitous autocorrelation in BOLD signals caused by physiological factors and unrelated to chunking. Each dataset contained 20 time series of *T* = 48 to match the 20-participant sample size for the empirical BOLD data. Emission probability *b*(·) was estimated from each time series separately and then averaged to serve as the initial for the Baum-Welch algorithm, which decoded one transition probability matrix *A* for each dataset.

To systematically test the effects of *d’* and *ρ* on decoding accuracy, each of the parameters varied from 0 to 0.9 in a step of 0.1, yielding 100 combinations. For each combination, 100 datasets were generated, each containing 20 time series. A decoding analysis was performed on each dataset following the same procedure above. The decoding accuracy was measured by the percentage of the 100 decoded matrices that followed the cardinal pattern of Eq. 2. Theoretically, the combinatorial probability of a random matrix following Eq. 2 is 1/84 (3 out of 9 positions could make 84 combinations) or 1.19%.

### Structural and functional MRI data acquisition

MRI data were acquired on a Siemens 3T Magnetom Prisma scanner with a 64-channel phased array head coil at the Center for Brain and Biomedical Imaging at The University of Delaware. Whole-head, high resolution structural images, including a T1-weighted, magnetizations-prepared rapid gradient-echo (MPRAGE) anatomical volume (TR=2500 ms, TE=2.9 ms, TI=1070 ms, flip angle=8.0°, voxel resolution=1.0 mm isotropic, FOV=256 × 256, 176 sagittal slices) and a T2-weighted anatomical volume (TR=3200 ms, TE=565 ms, flip angle=2.0°, voxel resolution=1.0 mm isotropic, FOV=256 × 256, 32 sagittal slices) were collected. Functional images were acquired using simultaneous multi-slice, T2*-weighted echo-planar imaging scans (TR=800 ms, TE=32 ms, flip angle=61°, FOV=21 cm, matrix=64 × 64, acceleration factor=6). We acquired 60 adjacent slices in an interleaved sequence with 2.5 mm slice thickness resulting in an in-plane resolution of 2.5 × 2.5 × 2.5 mm^3^.

### Image preprocessing

MRI data were preprocessed using the analysis pipeline of FreeSurfer (7.1.0) (Fischl, 2012) involving intensity normalization, registration to Talairach space, skull stripping, segmentation of white matter, tessellation of the white matter boundary, and automatic correction of topological defects (Fischl & Dale, 2000). The hippocampus mask was created in each subject’s native space using the utility provided by (Iglesias et al., 2015). BOLD preprocessing used a combination of SPM12 (Penny et al., 2011), FSL (5.0.9) (Smith et al., 2001), AFNI (Cox & Hyde, 1997) and FreeSurfer scripts implemented in Nipype 1.1.9 (Gorgolewski et al., 2011). Preprocessing steps involved dropping the 4 initial frames, despiking (AFNI), slice time correction (SPM SliceTiming), motion correction (SPM Realign), removal of linear and quadratic trends (FSL TSNR), and artifact detection based on movement or deviation in intensity (Nipype ArtifactDetect). Because of our subsequent autocorrelation and HMM analyses, no smoothing was applied to avoid distortion in the BOLD time series. Lastly, each subject’s T1 volume was co-registered to the functional space using boundary-based registration with 9 degrees of freedom (FreeSurfer bbregister).

### BOLD autocorrelation

BOLD time series were extracted from hippocampal voxels using the segmentation masks. Each data point followed immediately after each stimulus onset. For each voxel, lag-1 autocorrelation *r* was calculated following the formula:

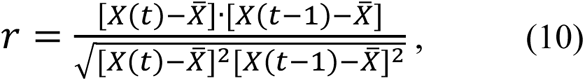

where *X*(*t*-1) = [*x*_1_, *x*_2_, …, *x_T_*_-1_] and *X*(*t*) = [*x*_2_, *x*_3_, …, *x_T_*_-1_] were BOLD time series of length *T*-1, shifted from each other by one time step, 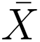 was the temporal mean of the full series [*x*_1_, *x*_2_, …, *x_T_*], and dot denoted inner product. For subject-averaged comparison of the hippocampal voxel histograms, the BOLD autocorrelation was resampled to the MNI152 T1 1mm template in FSL using FreeSurfer’s mri_robust_register algorithm. For all other analyses, *r* was extracted from each subject’s native space. When using the BOLD autocorrelation to estimate *ρ* in the chunking model, Eq. 10 became equivalent to Eq. 9, such that *X*(*t*-1) corresponded to o_t-1,i_, *X*(*t*) corresponded to o_t,j_, 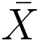 was calculated separately for *X*(*t*-1) (corresponding to 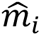) and *X*(*t*) (corresponding to 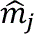). Emission probability *b*(·) was estimated in each subject separately and then averaged to serve as the prior for the Baum-Welch algorithm.

## Acknowledgements

This work was supported in part by National Institute on Deafness and Other Communication Disorders (NIDCD) Grant R21DC010576 to Z.Q. The data collection was conducted at the University of Delaware. The main analysis was conducted on the high-performance computing platform Big Red 3 and Carbonate, supported in part by Lilly Endowment, Inc. for the Indiana University Pervasive Technology Institute. We thank the staff at the Center for Biological and Brain Imaging at the University of Delaware for their critical support for our neuroimaging data collection. We thank Violet Kozloff and An Nguyen for their assistance in stimulus construction and programming for the experiment, Julie Schneider, Jennifer Legault, and Yi-Lun Weng for their contribution in data collection and organization, Anqi Hu for her assistance in behavioral data processing, and Marc Howard, Samantha Wood, Zoran Tiganj, Ho Ming Chow, Diane Chugani, and Frances Sayako Earle for helpful comments and discussions on the manuscript.

## FIG

**Supplemental Figure.**
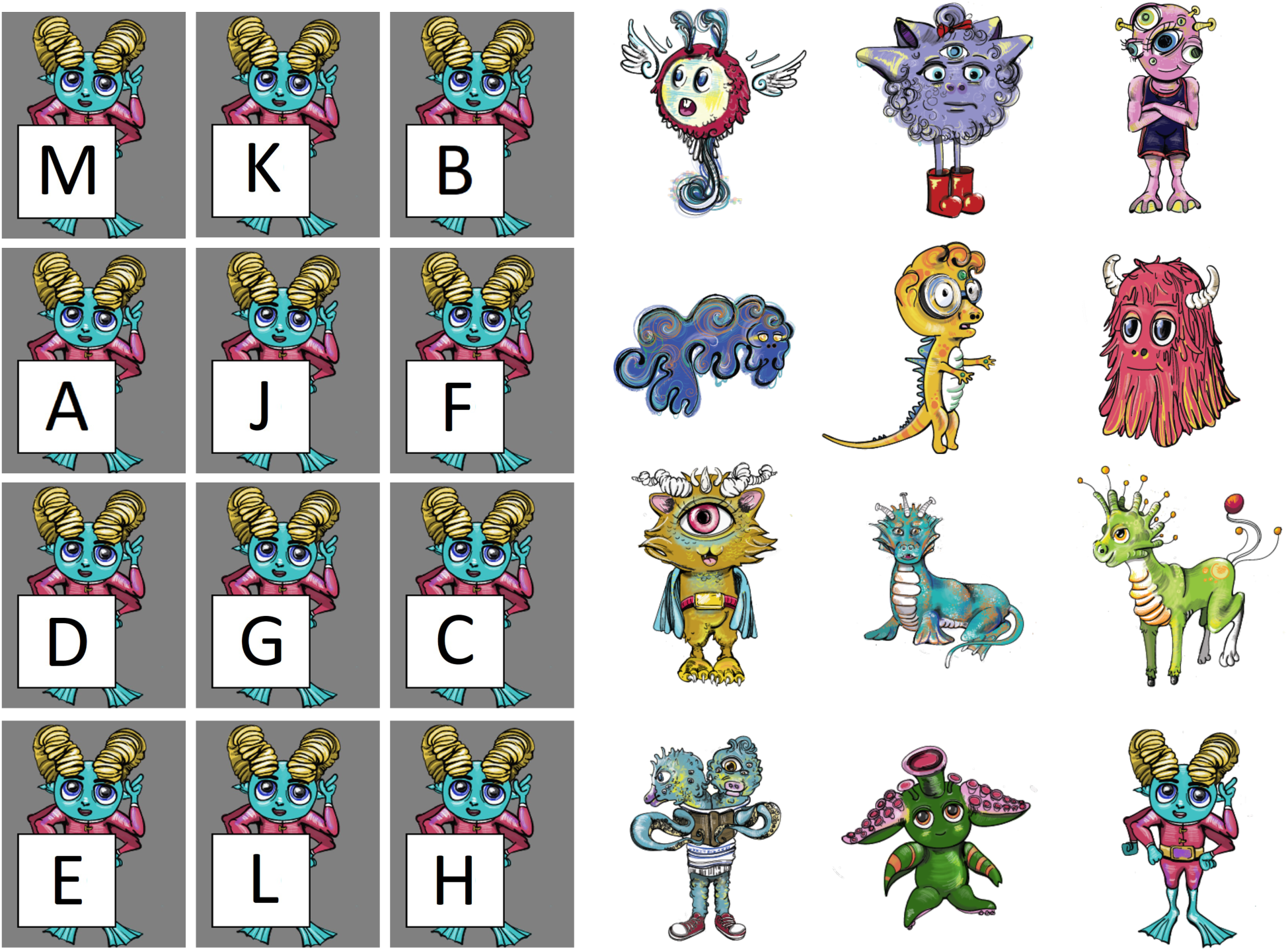
All the linguistic (left) and nonlinguistic (right) stimuli used in the task. For each type of stimuli, the three items on each row represent a triplet used in S-blocks.

## Footnotes

1 The exact nature of such generalization is complex: there is evidence for both modality– and stimulus-specific constraints on SL (see e.g., Frost et al., 2015, 2019, for reviews). As a first approximation, we here assume generalization across the perceptual domains of letters and hominoid figures within the visual modality.

2 Not exactly always because of the method’s probabilistic nature.

